# Time-series Multi-spectral Imaging in Soybean for Improving Biomass and Genomic Prediction Accuracy

**DOI:** 10.1101/2021.09.30.462675

**Authors:** Kengo Sakurai, Yusuke Toda, Hiromi Kajiya-Kanegae, Yoshihiro Ohmori, Yuji Yamasaki, Hirokazu Takahashi, Hideki Takanashi, Mai Tsuda, Hisashi Tsujimoto, Akito Kaga, Mikio Nakazono, Toru Fujiwara, Hiroyoshi Iwata

**Affiliations:** Department of Agricultural and Environmental Biology, University of Tokyo, Tokyo, Japan; Arid Land Research Center, Tottori University, Tottori, Japan; Graduate School of Bioagricultural Sciences, Nagoya University, Nagoya, Japan; Faculty of Life and Environmental Sciences, University of Tsukuba, Japan; Tsukuba Plant Innovation Research Center, University of Tsukuba, Tsukuba, Japan; Soybean and Field Crop Applied Genomics Research Unit, Institute of Crop Science, National Agriculture and Food Research Organization, Tsukuba, Japan

**Keywords:** multi-spectral (MS) imaging, time-series, above-ground biomass (AGB), genotypic values, phenotypic values, multi-trait (MT) genomic prediction, multi-kernel genomic prediction

## Abstract

Multi-spectral (MS) imaging enables the measurement of characteristics important for increasing the prediction accuracy of genotypic and phenotypic values for yield-related traits. In this study, we evaluated the potential application of temporal MS imaging for the prediction of above-ground biomass (AGB) and determined which developmental stages should be used for accurate prediction in soybean. Field experiments with 198 accessions of soybean were conducted with four different irrigation levels. Five vegetation indices (VIs) were calculated using MS images from soybean canopies from early to late growth stages. To predict the genotypic values of AGB, VIs at the different growth stages were used as secondary traits in a multi-trait genomic prediction. The accuracy of the prediction model increased starting at an early stage of growth (31 days after sowing). To predict phenotypic values of AGB, we employed multi-kernel genomic prediction. Consequently, the prediction accuracy of phenotypic values reached a maximum at a relatively early growth stage (38 days after sowing). Hence, the optimal timing for MS imaging may depend on the irrigation levels.

## 1. Introduction

Multi-spectral (MS) imaging is an effective method for assessing certain conditions in plants. Researchers have investigated the relationships between certain vegetation indices (VIs)—calculated based on spectral reflectance—and plant conditions [1][2][3][4][5][6][7]. MS image data and VIs have long been used in satellite remote sensing. Recently, small unmanned aerial vehicles (UAVs) equipped with MS cameras have been employed to collect high-resolution MS image data from plants. UAVs allow for the segmentation of pixels belonging to the subject (the plant) from those belonging to other objects (e.g., the background soil surface) [8][9][10]; thus, this method can sense certain conditions more accurately than satellite remote sensing[11]. With the development of these technologies, it has become possible to use MS image data and VIs to evaluate the state of a plant and predict future growth. In soybean (*Glycine max* L.), it has been reported that particular VIs from the growth stage can be used to predict grain yield. [12][13].

MS image data is also used with genome-wide marker data to predict certain traits (genomic prediction), as first demonstrated by [14]. The genetic correlation between MS image data and the target traits can be used to improve the prediction accuracy of genotypic values for said target traits. This approach is expected to improve selection accuracy for key breeding traits such as yield; thus facilitating genetic gain. The use of such models, incorporating VIs as secondary traits during the growth stage, have increased prediction accuracy for yield in maize and wheat compared to the standard genomic prediction model [15][16].

It is sensible to imagine that MS imaging will be used in combination with remote sensing, such as UAVs, and other -omics technologies to improve the efficiency and accuracy of genetic improvement in crops. In this case, the timing of measurements is critical, as the phenological growth stages for a given crop will affect image data [17][18][19], particularly soybean [20]. In soybean, previous studies have only performed MS imaging after the flowering stage [21][13]. There has been little to no evaluation of the potential for MS imaging from early to late growth stages in soybean. Moreover, the potential for MS imaging to predict genotype and phenotype for yield-related traits has not been investigated. MS imaging during the growth process will enable for the evaluation of changes in heritability and genetic and phenotypic correlations with target traits, and eventually to identify the appropriate measurement time for predicting a given target trait. MS image-based prediction of the genotypic values for a target trait may increase the accuracy of selection, improving the efficiency of breeding programs. Improved prediction of the phenotypic values for a target trait based on MS imaging would facilitate cultivation management and be a great help in farmers’ decision-making.

We observed above-ground biomass (AGB) as the target yield-related trait and collected MS images from early to late growth stages for 198 soybean accessions grown under different drought conditions. Drought stress generally causes severe damage to crop growth and ultimately lowers yield. Soybean, which serves as a major source of plant-derived protein and oil [22], exhibits a 40% reduction in yield due to drought on the global scale [23]. Clarifying the genotypic and phenotypic relationships between AGB and MS image data, we may be able to use MS imaging for genetic improvement of drought tolerance in soybean or efficient water-saving agronomic management.

We aimed to estimate the genetic correlations between VIs and AGB in different drought conditions over plant growth. Further, this work sought to predict the phenotypic values of AGB using the correlation between VIs and AGB. Lastly, we predicted the phenotypic values of AGB using the correlation between VIs and AGB across all treatment conditions; thus, evaluating whether macro-environmental variability (i.e., the differences in irrigation levels) is reflected in the VIs.

## 2. Materials and Methods

### 2.1. Experimental Setup and Data Collection

The accessions of soybean used, 198 in total, were obtained from the National Institute of Agrobiological Sciences (NIAS) gene bank. In 2019, all accessions were cultivated and evaluated at the Arid Land Research Center, at Tottori University in japan (35°32’N, 134°12’E, 14 m above sea level). The experimental field soil was sandy and retained high water permeability. On one ridge, two rows consisting of multiple micro plots were placed in parallel. Each micro plot contained four plants. The distances between two rows, two micro plots, and two plants were 50, 80, and 20 cm, respectively. Fertilizer (13, 6.0, 20, 11, 7.0 g m-2 of N, P, K, Mg, and Ca, respectively) was applied to the field prior to sowing. Sowing was performed on July 10th in 2019. Two to three seeds were sown at each position, after which germinated seedlings were thinned to one per position two weeks after sowing.

Four levels of watering treatment were employed—watering every day (control, C), no watering (drought, D), five days of watering followed by five days without (W5), and ten days of watering followed by ten days without (W10)(Figure S1). White mulching sheets (Tyvek, Dupond, US) were laid over the ridges to prevent rainwater infiltration into the soil and control soil drought levels. A watering tube was installed under the sheets between two rows in each ridge (Figure 1). Watering was done over five hours total daily (7:00–9:00, 12:00–14:00, 16:00–17:00) starting the day after seedling thinning for treatments C, W5, and W10. Soil moisture was measured using a soil moisture meter (TDR-341F, Fujiwara Seisakusho, Japan) at 48 sites in the field over 30 days (Figure 2). Data from several accessions were not used in the following analyses because of the unavailability of phenotypic records. The final dataset used contained 178 accessions in C, 183 in D, 188 in W5, and 184 in W10 (Table S1). For destructive AGB measurements at each plot, two plants located at the centre were taken from Sept 9 to 11 and the average was calculated. Days to flowering (DTF), which was defined as the date that 50% of plants flowered in each plot (Figure S2), was measured. To record DTF, we visited each plot every other day during the flowering period

**Figure 1.**
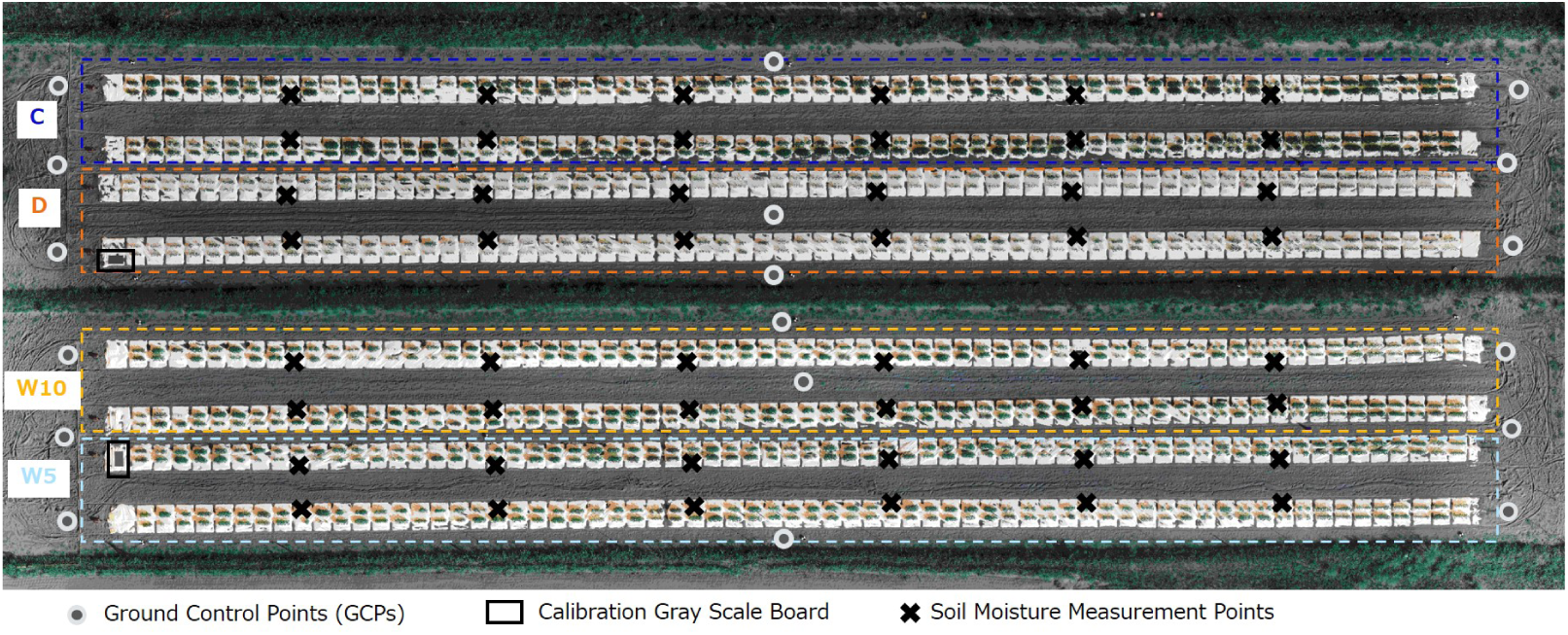
Experimental field overview and treatments; control (C), drought (D), five days of watering followed by five without (W5), and ten days of watering followed by ten without (W10).

**Figure 2.**
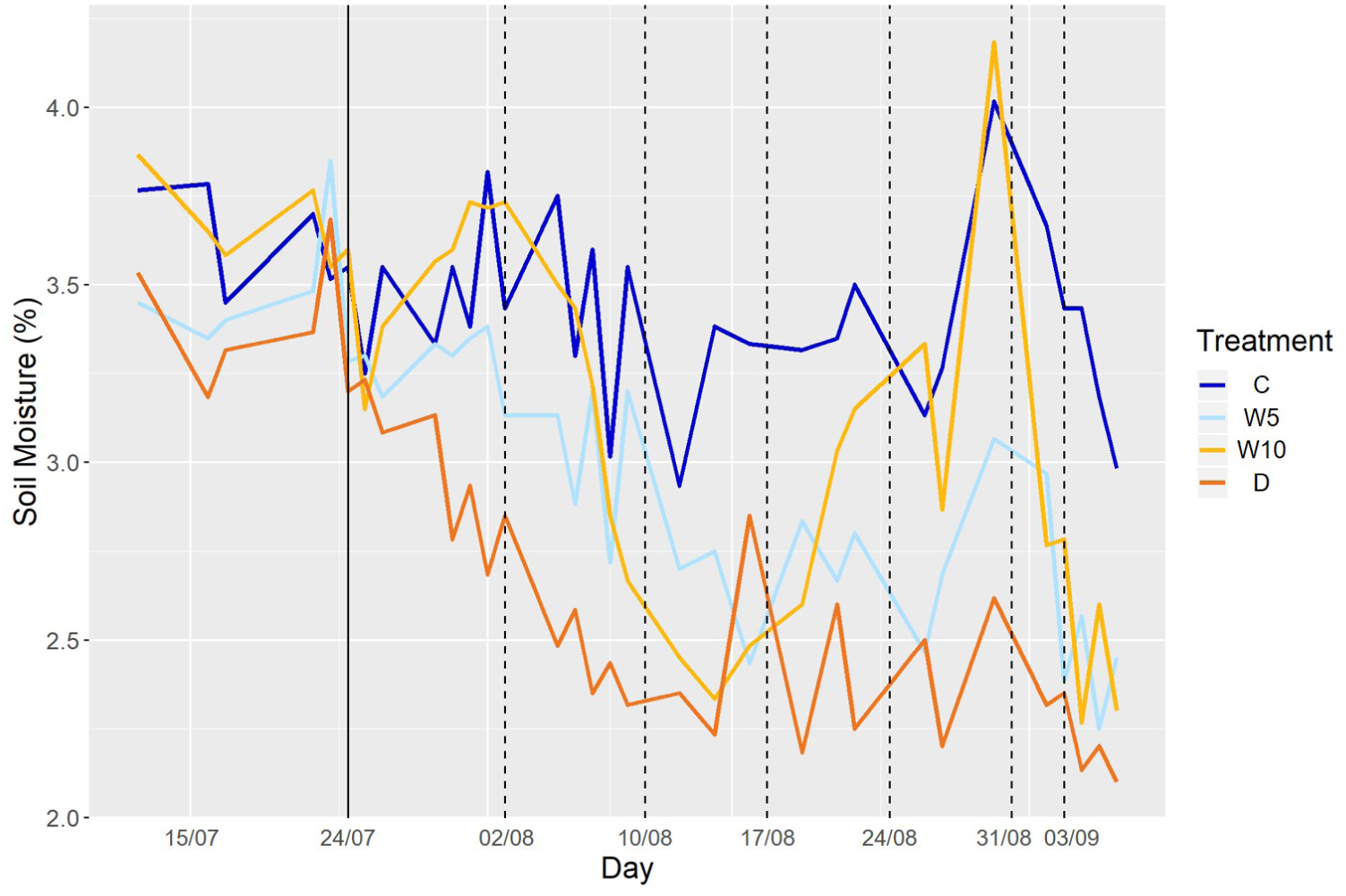
Soil moisture in each treatment. The solid line represents the date of thinning and dashed lines represent the dates for UAV measurements.

### 2.2. Multi-Spectral Data Collection and Processing

MS images (1920 × 1080 in resolution) were collected with a four-eye MS camera (Xacti, Japan) mounted on a UAV (DJI Matrice 100, DJI, China). The MS camera had four independent lens and sensors attached with different filters (MidOpt, USA), including two triple bandpass filters (TB475/550/850 and TB550/660/850), a red edge bandpass (Bi725), and a visible color bandpass (SP644). The spectral intensity of incident light was measured using a AS7265X DEMO KIT (ams, Austria) equipped with an 18-band MS sensor attached to the top of the UAV. MS imaging was calibrated using the spectral intensity of incident light. Images were collected six times (once a week) from August 2nd to September 3rd (Table 1). UAV flights were set to an altitude of 20 m, ensuring that the ground-level resolution was 1 cm/pixel, and scheduled for 12:00 under clear sky conditions daily. The ortho-mosaic images (OMIs) of four independent lens were obtained using Pix4Dmapper (Pix4D, Switzerland). Eighteen ground control points (GCPs) were set in the field. Using the geolocation information from the GCPs, 792 micro plots, each with four a maximum of plants, were segmented out of the OMIs. Plants were segmented out from an image of each micro plot to extract and analyze the MS image data only from the plants. NDVI differs significantly from the soil surface, to white mulching sheets, and plants (Figure S3); thus, plants were segmented with the NDVI threshold set to 0.15.

**Table 1.** Dates for MS imaging.

| Week | Date | Days after sowing (DAS) |
| --- | --- | --- |
| week 1 | Aug 02 | 23 DAS |
| week 2 | Aug 10 | 31 DAS |
| week 3 | Aug 17 | 38 DAS |
| week 4 | Aug 24 | 45 DAS |
| week 5 | Aug 31 | 52 DAS |
| week 6 | Sept 03 | 55 DAS |

### 2.3. Vegetation Index

Five types of VIs were used—the normalized difference vegetation index (NDVI) [1][2], green red vegetation index (GRVI) [3], normalized difference red-edge index (NDRE) [4], normalized difference index using red-edge and red spectral reflectance (NDI_red_) [6], and red-edge triangular vegetation index (RTVI) [7]. The VIs were calculated for each segmented plant pixel in each plot (Table 2). Each VI representing each plot was the median value of all segmented plant pixels in each plot.

**Table 2.**
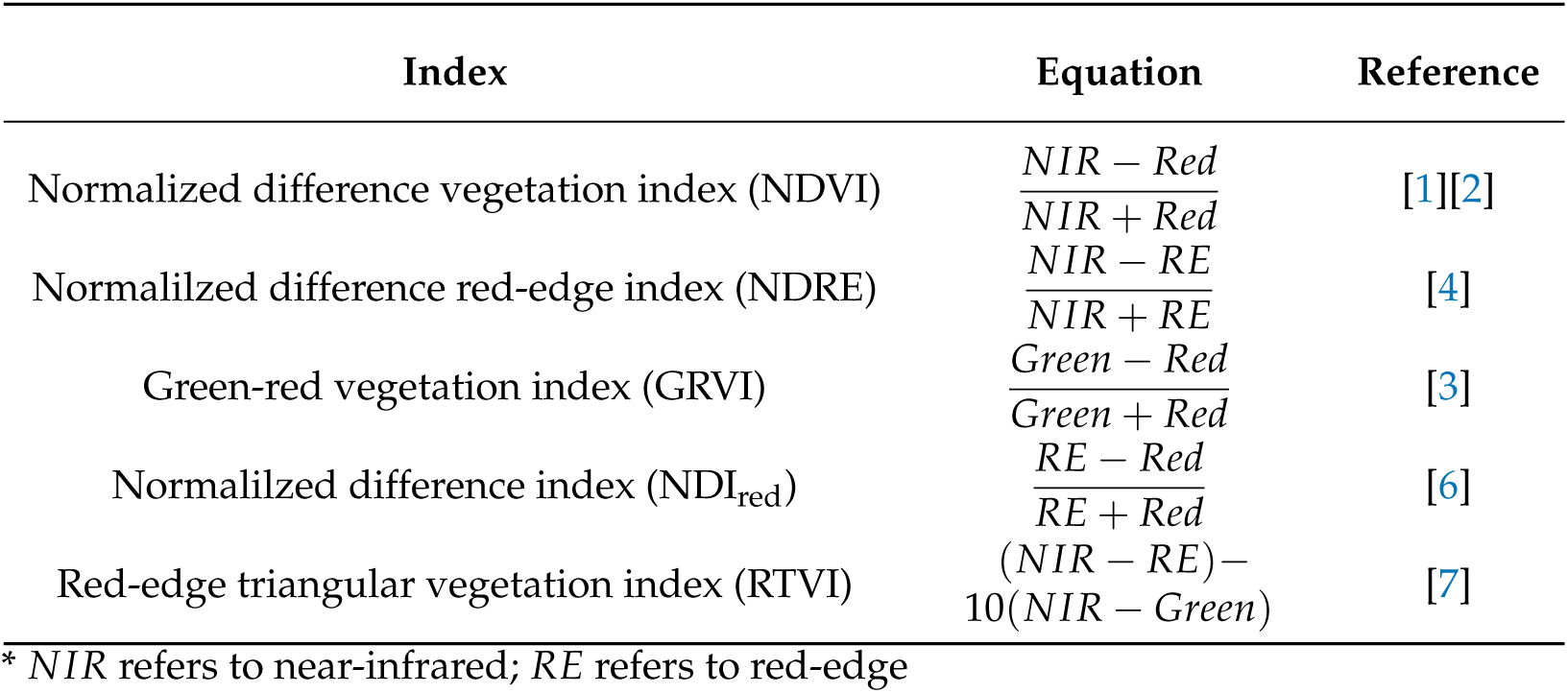
Vegetation Indexes (VIs) used in this study

### 2.4. Whole-genome SNP Data

Whole-genome sequence data for all accessions used were obtained [24]. All accessions were genotyped using 4,776,813 single nucleotide polymorphisms (SNPs) filtered using certain criteria. The SNPs of heterozygous and if more than 95% of the individuals had missing data for a SNP, they were removed. The markers were also filtered for a minor allele frequency (MAF) less than 0.025, and missing data were imputed based on the mean, after which they were filtered again for a MAF less than 0.05. Finally, the linkage disequilibrium (LD) was computed only for SNP pairs for which the distance was less than 25,000 base pairs (bp) and SNPs were selected at LD below 0.8, resulting in a total of 173,583 SNP markers. This thinning with LD was performed using the ‘LD.thin’ function in the ‘gaston’ package in R version 1.5.7 [25]. The additive numerator relationship matrix **G** was estimated using the ‘calcGRM’ function in the ‘RAINBOWR’ package in R version 0.1.25 [26].

### 2.5. Genomic Heritability

Genomic heritability [27] was estimated to determine the level of genetic control of VIs at each growth stage and each treatment. Genomic heritability is the proportion of phenotypic variance explained by genome-wide SNPs [28][29]. Genomic heritability for each of VIs and for AGB was calculated following *G*:

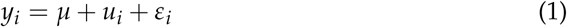

where *y*_*i*_ is the phenotypic value for each VIs or AGB for genotype *i* (*i* = 1, …, *n*), *μ* the overall mean, *u*_*i*_ the genetic random effect, and *ε*_*i*_ the residual random effect. Vector **u** = (*u*_1_, …, *u*_*n*_)^*T*^ follows the multi-variate normal distribution, 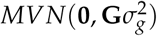, where **G** is the additive genomic relationship matrix and 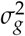 is the additive genetic variance. Vector ***ε*** = (*ε*_1_, …, *ε*_*n*_)^*T*^ follows the multi-variate normal distribution 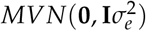, where **I** is the identical matrix and 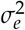 is the residual variance. Genomic heritability was calculated as: 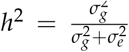 The model was implemented using the ‘EMM.cpp’ function in the ‘RAINBOWR’ package in R version 0.1.25 [26].

### 2.6. Multi-trait Model

For each combination of growth stage (week) and treatment, the multi-trait (MT) model below was built using a Bayesian multivariate Gaussian model [30][31] to estimate genetic and residual correlations between the VIs and AGB. This model was also used to predict the genotypic values of AGB.

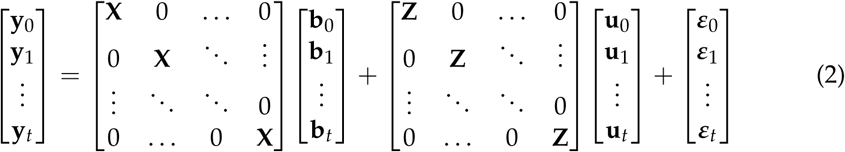

where *t* is the number of VIs (*t* = 5), **y**_0_ and **y**_*k*_ (*k* = 1, …, *t*) the vectors of *n* × 1 length phenotypic trait value of AGB and VI, **X** the fixed effect design matrix (which is the same for all traits), **b**_*k*_ (*k* = 0, …, *t*) the vector of fixed effects of flowering for trait *t*, **Z** the random effect design matrix (the same for all traits), **u**_*k*_ the vector of genetic random effects of trait *k*, and ***ε***_*k*_ the vector of residual random effects of trait *k*. The vector. 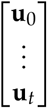 follows a multi-variate normal distribution *MVN*(**0, G** ⊗ **Σ**), where **G** is the additive genomic relationship matrix and **Σ** is the genetic variance-covariance matrix across traits. The vector 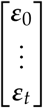 follows the multi-variate normal distribution *MVN*(**0, I** ⊗ **R**), where **I** is the identical matrix, **R** the residual variance-covariance matrix across traits, and **I** the identical matrix. **Σ** and **R** were estimated as a *t* × *t* unstructured matrix. This model was built using the ‘MTM’ R package version 1.0.0 [32].

### 2.7. Single/Multiple Kernel Models within Each Treatment

To predict the phenotypic values of AGB for each treatment, we built four kernel regression models—(1) with the additive genomic relationship (**G**) kernel, (2) with the VIs relationship (**V**_*each*_) kernel, (3) with **G** and **V**_*each*_ kernels, and (4) with **G** and the field heterogeneity relationship kernel (**FH**) (Table 3).

**Table 3.** Single/Multiple kernel models within each treatment.

| Model Name | Kernel | Equation | Index |
| --- | --- | --- | --- |
| G model | **G** | $y_{il} = b_l + u_i + \varepsilon_i$ | Equation S1 |
| **V<sub>each</sub>** model | **V<sub>each</sub>** | $y_{il} = b_l + v_i + \varepsilon_i$ | Equation S2 |
| **G** + **V<sub>each</sub>** model | **G, V<sub>each</sub>** | $y_{il} = b_l + u_i + v_i + \varepsilon_i$ | Equation S3 |
| **G** + *Field* model | **G, FH** | $y_{il} = b_l + u_i + f_i + \varepsilon_i$ | Equation S4 |

\* **G**: the additive genomic relationship kernel, **V<sub>each</sub>**: the VIs relationship kernel for each treatment, **FH**: the field heterogeneity relationship kernel. $y_{il}$ : the phenotypic value of AGB for genotype $i$ ( $i = 1, \dots, n$ ), $b_l$ : the fixed effect of flowering ( $l = 1, 2$ ), $u_i$ : the genetic random effect, $v_i$ : the random effect of the VIs, $f_i$ : the random effect reflecting the among-plot distances, and $\varepsilon_i$ : the residual random effect.

The single-kernel model with **G** (1) compares the utility of a VI-based model to a standard genome-wide marker-based model. The single-kernel model with **V**_*each*_ (2) evaluates the usefulness of VIs in predicting the phenotypic values of AGB within each irrigation level. This model can be used to predict the phenotypic values of AGB based on MS image data at a low cost, without genome-wide marker data and destructive sampling. The multi-kernel model with **G** and **V**_*each*_ (3) was employed to compare whether the combination of genome-wide marker and VIs data would increase the prediction accuracy. The multi-kernel model with **G** and **FH** (4) aimed to confirm whether the field heterogeneity affected the phenotypic values of AGB within each treatment. When this model exhibits no or little increment of the prediction accuracy compared to the **G** single-kernel model, then the location effect of the plots (i.e., distance dependent among-plot heterogeneity) can be assumed to be small in this experimental field.

The details of these three types of kernels and four types of single/multiple kernel models are described in the *Appendix S1*.

### 2.8. Single/Multiple Kernel Models over All Treatments

To evaluate whether macro-environmental variability, i.e., the differences in irrigation levels, is reflected in the VIs, we built models using data of all treatments. We built four kernel regression models: (1) A model with the additive genomic relationship kernel (**G**), fixed effects of treatment (E), and genotype-by-environment interaction (*G* × *E*) effect, (2) A model with **G**, E, and VIs relationship kernel (**V**_*all*_), (3) A model with the **G** and **V**_*all*_, (4) A model with **V**_*all*_ (Table 4).

**Table 4.** Single/Multiple kernel models over all treatments.

| Model Name | Kernel | Equation | Index |
| --- | --- | --- | --- |
| $G + E + G \times E$ model | **G, GE** | $y_{ijl} = E_j + b_l + u_i + ge_{ij} + \varepsilon_{ij}$ | Equation S5 |
| $G + E + V_{all}$ model | **G, $V_{all}$** | $y_{ijl} = E_j + b_l + u_i + v_{ij} + \varepsilon_{ij}$ | Equation S6 |
| $G + V_{all}$ model | **G, $V_{all}$** | $y_{ijl} = b_l + u_i + v_{ij} + \varepsilon_{ij}$ | Equation S7 |
| $V_{all}$ model | **$V_{all}$** | $y_{ijl} = b_l + v_{ij} + \varepsilon_{ij}$ | Equation S8 |

\* **G**: the additive genomic relationship kernel, **GE**: genotype-by-treatment interaction kernel, **$V_{all}$** : the VIs relationship kernel over all treatment. $y_{ijl}$ : the phenotypic trait value of AGB for genotype $i$ ( $i = 1, \dots, n$ ) in treatment $j$ ( $j = 1, \dots, m$ ), $E_j$ : the fixed effect for treatment, $b_l$ : the fixed effect of flowering ( $l = 1, 2$ ), $u_i$ : the genetic random effect, $ge_{ij}$ : the genotype-by-environment interaction random effect, $v_{ij}$ : the random effect of the VIs, and $\varepsilon_{ij}$ : the residual random effect.

The aim of each of the four models is as follows. The multi-kernel model with **G**,E, and *G × E* (1 in the previous paragraph) aimed to compare the utility of a VI-based model to a standard genome-wide marker data and treatment information-based model. The multi-kernel model with **G**, E, and **V**_*all*_ (2 in the previous paragraph) aimed to evaluate the usefulness of VIs in modelling the *G × E*. When this model outperformed the prediction accuracy of the **G**, E, and *G × E* multi-kernel model, then the MS image data may be a useful phenotype for modelling *G × E*. The multi-kernel model with **G** and **V**_*all*_ (3 in the previous paragraph) was aimed to evaluate whether the differences in irrigation levels is reflected on Vis. This model can be used in a large field, drought levels can be largely complex and uneven. The single-kernel model with **V**_*all*_ was aimed to evaluate the usefulness of VIs for predicting the phenotypic values of AGB over all irrigation levels. This model can be used for predicting phenotypic values of AGB based on MS image data at a low cost (without genome-wide marker data and destructive sampling).

The details of these four models are described in the *Appendix S1*. All single/multiple kernel models were built using ‘EM3.cpp’ function in the ‘RAINBOWR’ package in R version 0.1.25 [26].

### 2.9. Evaluation of Prediction Accuracy

The prediction accuracy of each model was evaluated using five-fold cross-validations with 10 replications. The accuracy of the multi-trait model was evaluated by using AGB only in the training set, while the phenotypic data from the VIs were used in both the training and test sets. For prediction pooled all treatment, the genotype used for the training data was not used for the test data. The distribution of AGB values in each treatment did not follow a normal distribution; thus, the genomic heritability of AGB was calculated after taking the logarithm—ensuring that the error term follows a normal distribution. The MT and single/multiple kernel models were built using these log values for AGB. The correlations between the observed and predicted values were calculated by taking the exponential of the predicted values.

## 3. Results

### 3.1. Heritability and Phenotypic and Genetic Correlation of AGB and VIs Across Four Drought Treatment Levels

Average AGB descended from treatment C to D (Table 5). Similarly, the genomic heritability of AGB was the highest in treatment C and lowest in treatment D. This trend was also observed for the VIs (Figure 3a). For VIs, we evaluated the genomic heritability not only for the different drought treatments, but also for the different growth stages. The genomic heritability of the VIs increased in the later growth stages, with the exception of treatment D. The results suggest that AGB and VIs under severe drought conditions were more strongly affected by field heterogeneity compared to better-irrigated conditions. Next, we focused on the phenotypic correlation between AGB and the VIs. The correlation between AGB and each individual VI was similar between the different drought conditions (Figure 3b). The sign of the correlation in two VIs (GRVI and NDI) with AGB changed during growth stages in treatments W5 and D, while three VIs (NDVI, NDRE, and RTVI) showed positive correlations with AGB across all treatments. The correlation between AGB and VIs was higher in treatments C and D compared to W5 and W10, especially in the later growth stages (at week 5 and 6).

**Table 5.** Mean AGB (with standard error) and genomic heritability in each drought treatment

| treatment | C | W5 | W10 | D |
| --- | --- | --- | --- | --- |
| Mean (g) ± SE | 73.29 ± 2.33 | 62.05 ± 1.67 | 53.47 ± 1.48 | 16.85 ± 0.63 |
| genomic heritability | 0.48 | 0.41 | 0.44 | 0.26 |

**Figure 3.**
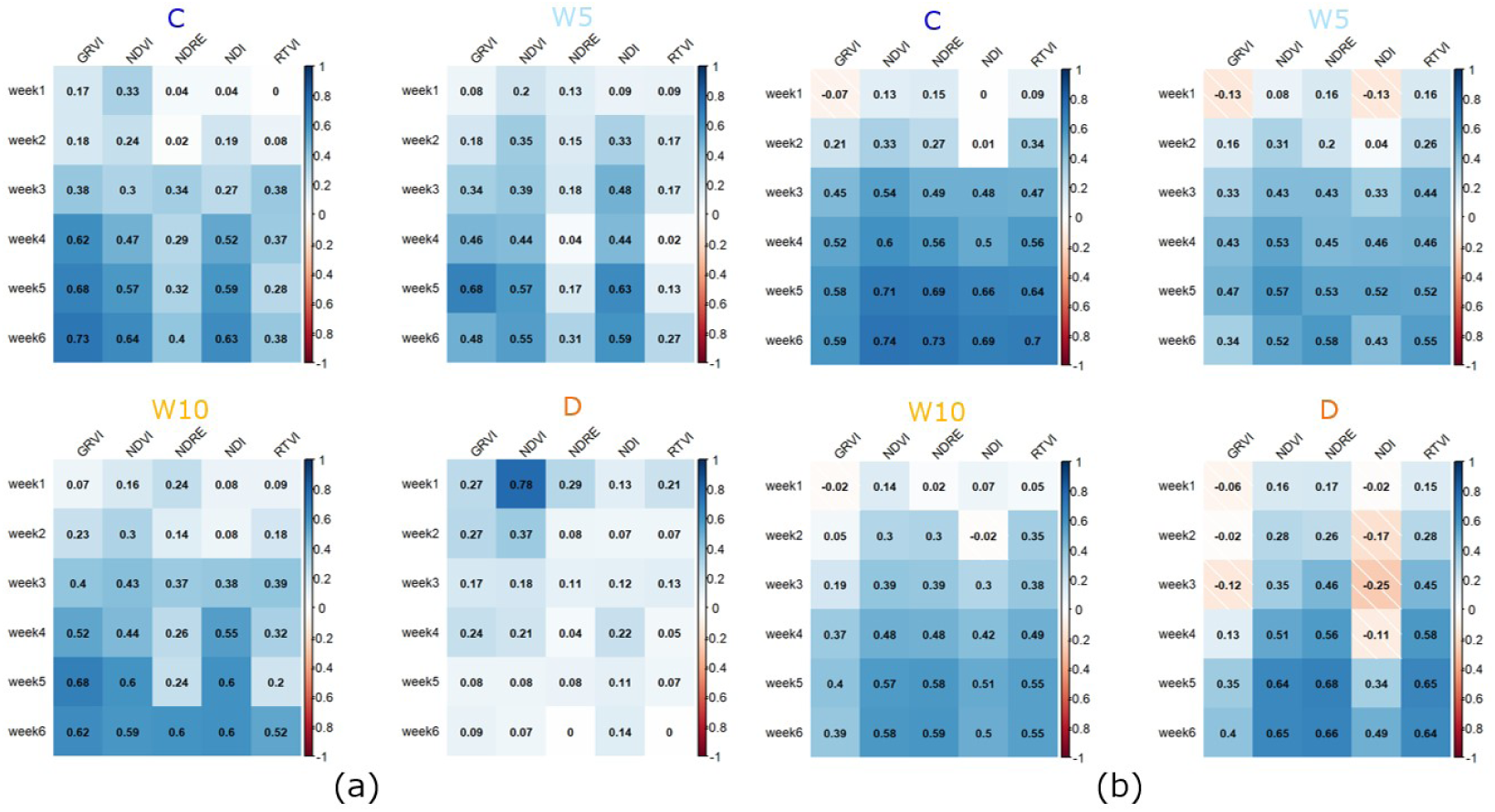
Genomic heritability and phenotypic correlation. (**a**) Genomic heritability for the VIs in each treatment. (**b**) Phenotypic correlation between AGB and VIs in each treatment.

### 3.2. Genetic Correlations Following the MT model

Genetic correlation was derived from the among-trait genetic variance-covariance matrix (**Σ**) estimated following the MT model (Equation (2)). In the 1st week, the genetic correlation under each treatment was low—C, −0.04; W5, −0.03; W10, 0.01, and D, −0.04 on average. In the 6th week, the genetic correlations increased to 0.75, 0.76, and 0.75 for treatment C, W5, and W10, respectively. Under treatment C, W5, W10 in the VIs with the highest correlation, whereas the genetic correlation increased only up to 0.37 with treatment D, in the VIs with the highest correlation (Figure 4). The genetic correlation between the VIs and AGB in treatment C, W5, and W10 increased drastically from the week 4-5, with an average increase of 240%. Additionally, GRVI and NDI showed a high positive genetic correlation at the late growth stages (week 5 and 6) in treatments C, W5, and W10, whereas a low and negative genetic correlation was observed for treatment D. In contrast, NDVI, NDRE, and RTVI showed positive genetic correlation with AGB in all drought treatment levels except the early growth stages (week 1 and 2). Overall, these results indicate that genetic correlations between VIs and AGB varied during growth and between drought treatment conditions.

**Figure 4.**
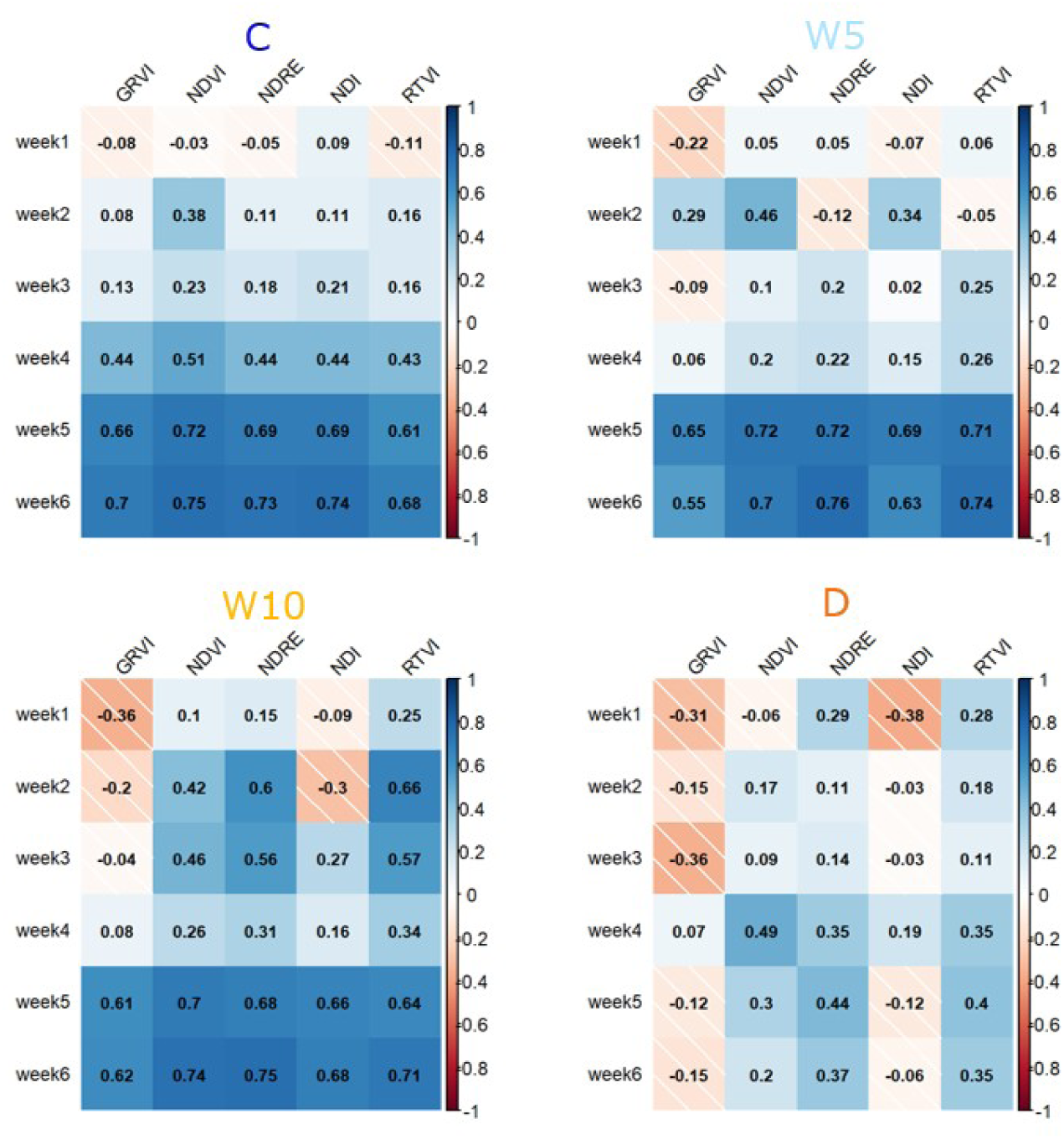
Genetic correlations between AGB and the VIs.

### 3.3. Prediction Accuracy of AGB using the MT model

The prediction accuracy of the genotypic values for AGB with models G and MT was evaluated using five-fold cross-validation (Figure 5). The prediction accuracy of model G changed by the fixed effect of flowering and non-flowering. First, we compared the prediction accuracy of models G and MT. The mean accuracy of model G for treatments C, W5, W10, and D was 0.39, 0.28, 0.32, and 0.18, respectively. Model MT was able to predict AGB more accurately than model G for all drought treatment conditions. At the 2nd week, the prediction accuracy of the model MT was already 32, 58, 45, and 55% higher than that of model G, for all treatments. The prediction accuracy of model MT increased significantly from the 1st to the 3rd week, averaging 70% in all treatments, but did not change much from the 3rd week on—averaging 6% for all treatments.

**Figure 5.**
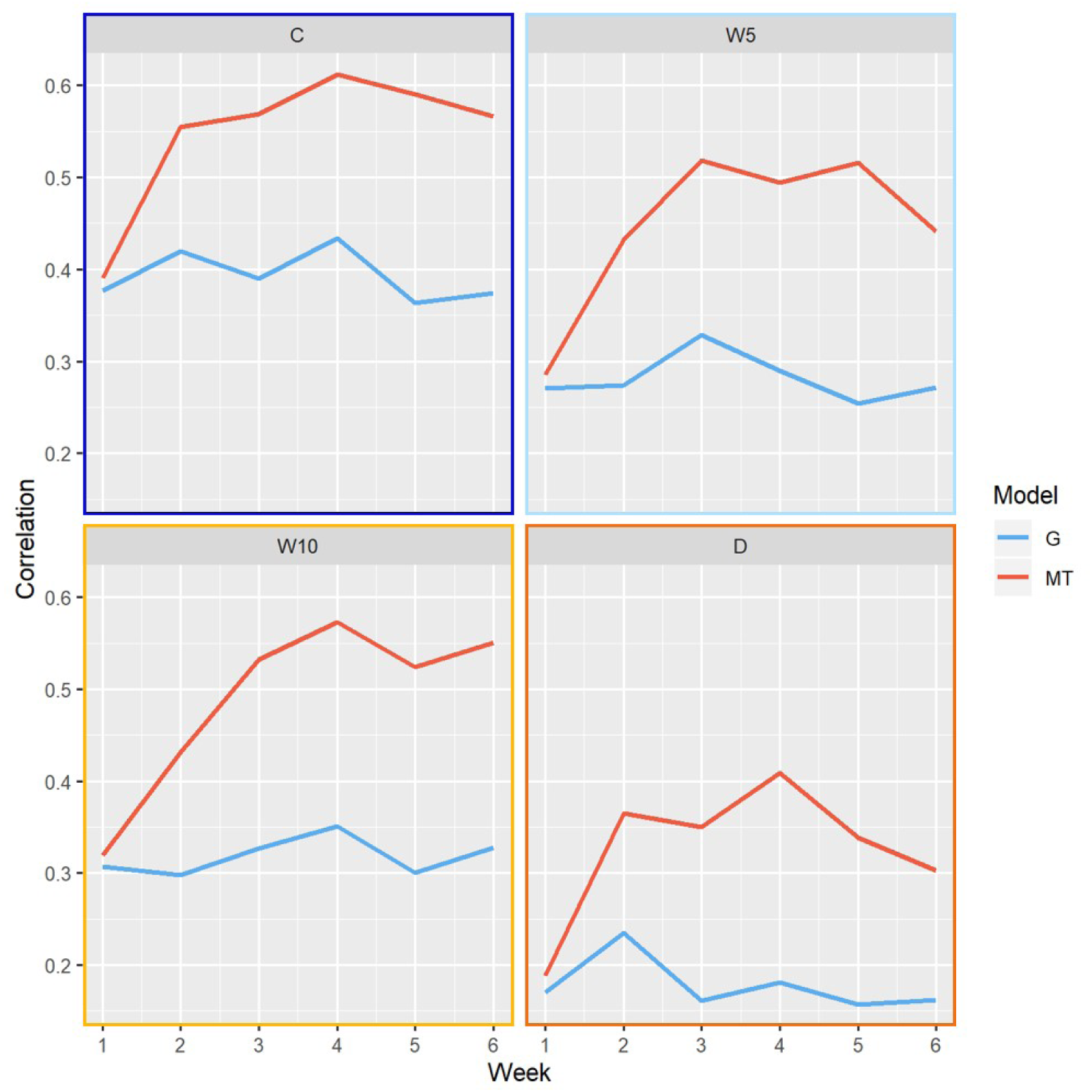
AGB prediction from models G and MT. Prediction accuracies were estimated as the correlation between observed and predicted values.

### 3.4. Prediction Accuracy of AGB with Single/Multiple Kernel Regression Models

Three types of single/multiple kernel regression models (*V*_*each*_, *G* + *V*_*each*_, and *G* + *Field* models) predicted the phenotypic values of AGB. Figure 6 shows the prediction accuracy of model *G* (Equation S1), *V*_*each*_ (Equation S2), and *G* + *V*_*each*_ (Equation S3). Model *G* + *Field* (Equation S4) hardly increased the prediction accuracy of model *G* (Figure 6). Except for the 1st week, the prediction accuracy of model *V*_*each*_ was higher than that of model *G*. In the 2nd week, the prediction accuracy of model *G* + *V*_*each*_ was 52% higher than that of model *G* for all treatments. For model *V*_*each*_, however, the prediction accuracy in treatments C, W5, and W10 reached its maximum in week 2 or 3 The prediction accuracy in treatment D gradually increased over time. Further, the maximum prediction accuracy values for model *G* + *V*_*each*_ were 0.67, 0.77, 0.58 and 0.60 in treatments C, D, W5, and W10, respectively—with the C and D being 22% higher than W5 and W10 on average.

**Figure 6.**
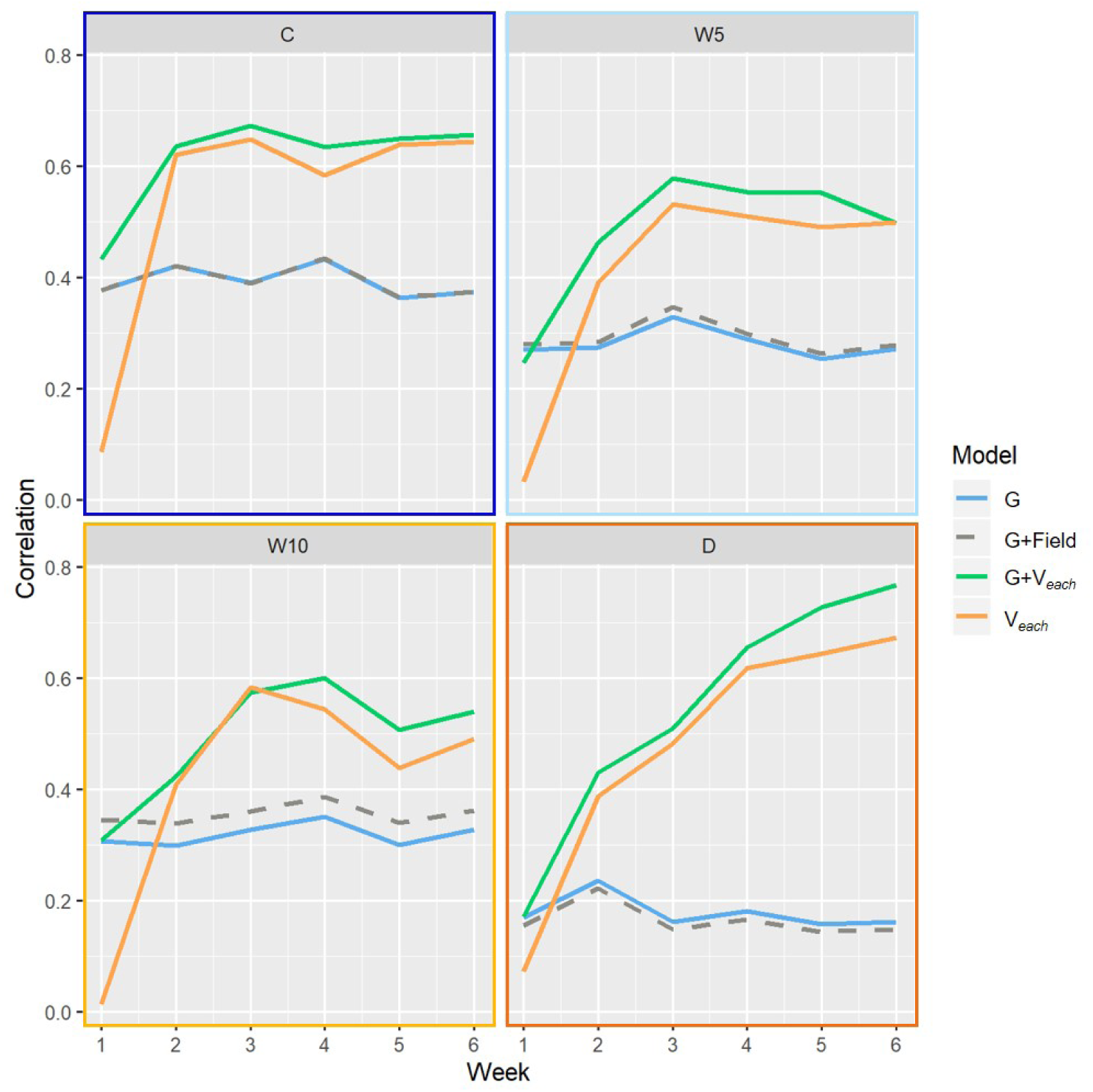
AGB prediction from single/multiple kernel regression models. The prediction accuracy was estimated as the correlation between observed and predicted values.

### 3.5. Prediction Accuracy for AGB Across All Drought Treatment Conditions

Four types of single/multiple kernel regression models (*G* + *E* + *G* × *E, G* + *E* + *V*_*all*_, *G* + *V*_*all*_, and *V*_*all*_ model) were used to predict the phenotypic values of AGB over all drought treatment conditions (Figure 7). The prediction accuracy of model *G* + *E* + *V*_*all*_ was always higher (by 10%, on average) than that of model *G* + *E* + *G × E*. Additionally, models *G* + *V*_*all*_ and *V*_*all*_ outperformed model *G* + *E* + *G* × *E* from week 4-6. These result suggest that the MS image data provided details on the plant growth conditions that were not captured by differences in the drought level or genomic information. The prediction accuracy of the three models (*G* + *E* + *V*_*all*_, *G* + *V*_*all*_, and *V*_*all*_) using MS image data increased significantly from week 1-4, after which it did not change much.

**Figure 7.**
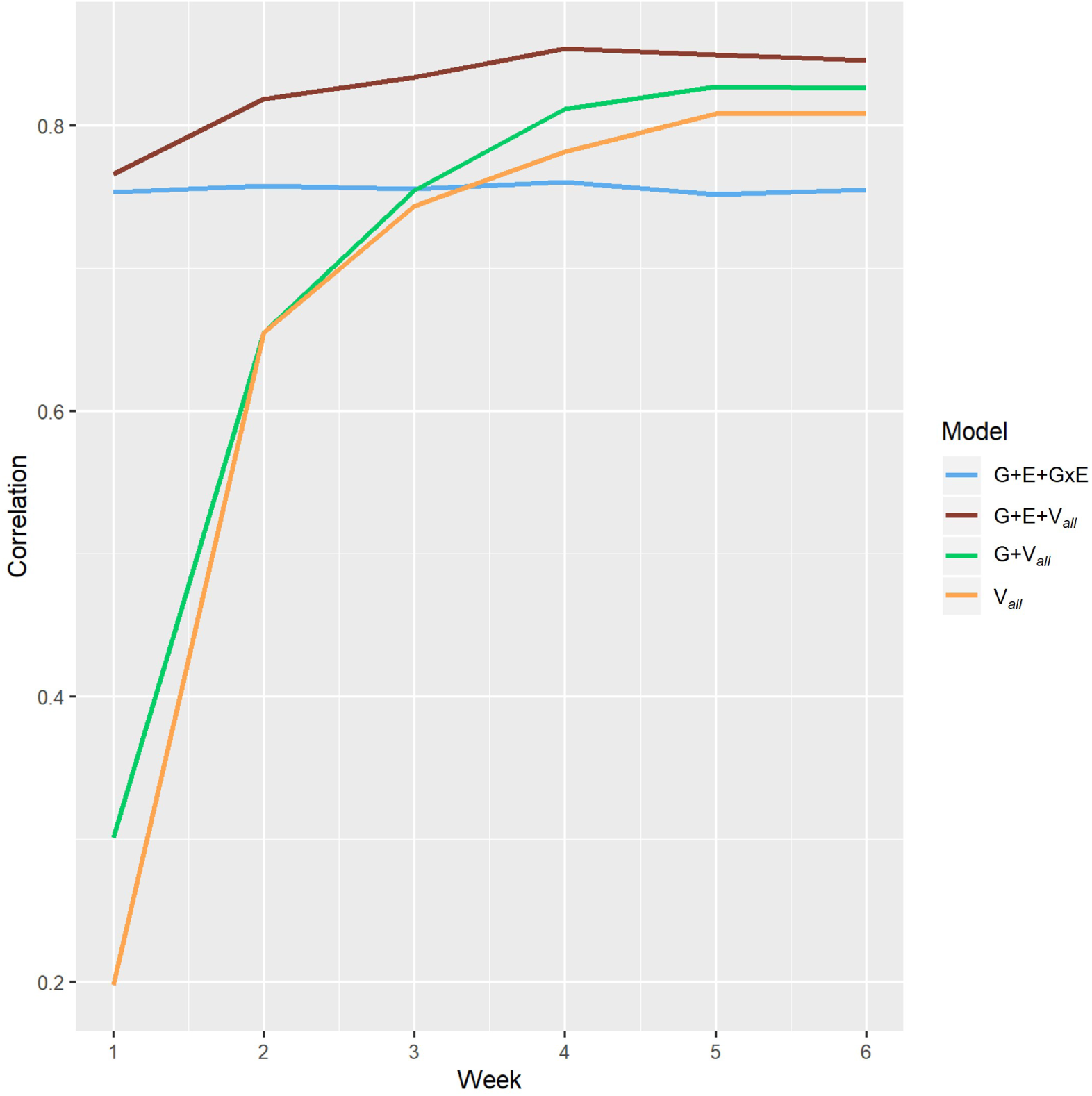
AGB prediction from single/multiple kernel regression models across all treatments. Prediction accuracy was estimated as the correlation between observed and predicted values.

## 4. Discussion

### 4.1. Heritability of the VIs

Overall, we found that VI genomic heritability increased with growth (Figure 3a). This is consistent with previous work [20] reporting that the genomic heritability of VIs was higher in the later growth stage in soybean. During the early growth stage, when the plants are smaller, reflectance from the ground could impact the MS data acquired [33][34]. However, in our treatment D, heritability of the VIs did not exhibit this increase. Soil moisture may have varied between plots, particularly in treatment D, as it is difficult to control the environmental conditions in the field. Further, drought is known to inhibit growth in soybean [23]. Under the severe drought conditions of treatment D, field heterogeneity in soil moisture is expected to have a larger impact on plant growth than the other treatments. It was reported that differences in soil moisture affect spectral reflectance in soybean [35][36]. Further, plant size in treatment D was significantly less compared to the other treatments, making the accurate segmentation of the plants difficult; thus, the background spectra could have introduced noise into the data.

### 4.2. Phenotypic and Genetic Correlations

The phenotypic correlation between AGB and three VIs (NDVI, NDRE, and RTVI) was larger in later growth stages, for all treatments conditions (Figure 3b). This suggests that the three VIs would serve as good indicators for AGB, no matter the irrigation conditions. However, a genetic correlation between AGB and the VIs, as estimated using the MT model, showed a clear difference between treatment D and the others (Figure 4). In treatment D, the phenotypic correlation between AGB and the VIs increased to 0.9 following growth, whereas genetic correlations increased up to 0.49. This suggests that the high phenotypic correlation is mainly due to the high genetic correlation under irrigated conditions, while the high phenotypic correlation is due to the high residual correlation under drought conditions. Hence, under the drought condition, VIs are able to reflect more of the micro-environmental variations resulting from field heterogeneity than the genetic variations from the different soybean accessions.

### 4.3. MT Model Prediction

The prediction accuracy for the genotypic values of AGB was improved in model MT compared with that in model G. To predict a target trait with low heritability, such as yield or biomass, it would be useful to incorporate secondary traits that have high heritability and high genetic correlation to the target trait [37].

Spectral reflectance in soybean is known to be influenced by the maturity of the plant [21]. It has been reported that spectral reflectance measurements also capture physiological parameters associated with relative maturity in wheat [38]. Therefore, wheat grain yield corrected based on the variations in days to heading reduced the advantage of model MT using NDVI as a secondary trait over model G [14]. In our study, the factors of flowering and non-flowering were included in models as fixed effects, though the advantage of model MT over G did not reduce in soybean.

When predicting the genotypic values of AGB, the accuracy of model MT began increasing early—31 or 38 days after sowing—no matter the treatment. The use of MS image data from the early growth stages allowed us to predict the genotypic values of AGB with higher accuracy than that of genomic information alone. Early growth stage data could allow breeders to cull genotypes with low predicted genotypic values before harvest, which would save time and labor. Thus, MT imaging is expected to improve the streamline and accuracy of selection in AGB and eventually yield.

### 4.4. The Single/multiple Kernel Regression Models

In the single/multiple kernel prediction, *G* + *V*_*each*_ and *V*_*each*_ outperformed *G* in all conditions, except in the 1st week (Figure 6). The models using kernel **V**_*each*_ increased the prediction accuracy of AGB compared to model *G* because they could leverage the phenotypic correlations between the VIs and AGB.

It has been reported that the initial seed-filling stage (R5) was the best development stage to predict soybean yield using MS image data [21][13]. However, in our study, the prediction accuracy of phenotypic values in *V*_*each*_ increased over time in treatment D. For the other conditions, the prediction accuracy reached a maximum at 38 days after sowing (week 3). It may be that MS data should be collected at different times, depending on the drought conditions of the field.

We expected that the growth environment would differ depending on the field heterogeneity in each condition; thus, a model with kernels **G** and **FH** (*G* + *Field* model) was also employed. However, the prediction accuracy was almost the same as that of model *G* (Figure 6)—suggesting that the VIs captured not only the field heterogeneity explained by the proximity between plots, but also the micro-environmental differences which could not be explained by kernel **FH**.

### 4.5. Prediction over All Treatment Levels

From the early growth stages to 38 days after sowing (week 3), the prediction accuracy of model *G* + *V*_*all*_ was lower than that of *G* + *E* + *G × E*, although the difference in soil moisture content was already observed between different treatments (Figure 2). This suggests the difficulty in the identification of drought level at which soybean plants grow with MS imaging occurs in an early growth stage.

The prediction accuracy of model *G* + *E* + *V*_*all*_ was 10% higher than that of *G* + *E* + *G × E* on average, indicating that the VIs can capture important differences in plant growth despite field heterogeneity between different treatment conditions. It was also reported that model *G* + *E* + *V*_*all*_ has higher prediction accuracy compared to *G* + *E* + *G × E* for wheat yield prediction [38]. In this study, *G* + *V*_*all*_ and *V*_*all*_ outperformed *G* + *E* + *G × E* in prediction accuracy after week 4. This indicates that MS imaging can capture not only micro-environmental variability, but also macro-environmental variability, i.e., the differences in irrigation levels.

## 5. Conclusions

We collected MS image data from the early to late growth stages of soybean and estimated the genetic and phenotypic correlations between VIs and the yield-related trait AGB. The heritability of VIs and the genetic correlation between the VIs and AGB were significantly different between different drought levels. The heritability and the genetic correlation increased over time, except under severe drought conditions. However, in model MT, the prediction accuracy of genotypic values in AGB increased at relatively early growth stages (31 or 38 days after sowing) in all drought treatments. This indicates that MS imaging would be useful for breeders in the selection of promising lines or plants. Early measurement of secondary traits will allow breeders to cull the unwanted plants before harvest, saving effort on time and labor. Furthermore, early selection using predicted genotypic values may contribute to the acceleration of breeding cycles.

The usefulness of MS imaging for cultivation management was also evaluated. Phenotypic values for AGB were predicted by kernel regression using the phenotypic correlations between the VIs and AGB. The prediction accuracy of model *G* + *V*_*each*_ (using VIs and genome-wide marker data) was higher than that of model *G* (using only genome-wide marker data) early in growth, no matter the drought conditions. This indicates that VIs were able to capture the environmental-variation in the phenotypic value of AGB, which cannot be captured by genome-wide marker data alone. Furthermore, the prediction accuracy of model *G* + *V*_*all*_ was higher than that of *G* + *E* + *G* × *E* across all treatments. This indicates that VIs can capture the variations in AGB caused by the macro-environment, i.e., different drought levels, in the later growth stages. In a large field, drought levels can be largely complex and uneven, which makes cultivation management difficult. In such fields, the estimation of drought levels and growth prediction using MS imaging in the later growth stages may be of great help for decision making in cultivation management.

## Supplementary Materials

The following are available online at https://www.mdpi.com/article/10.3390/1010000/s1, Figure S1: The cycle of irrigation and the timing of each management and measurement, Figure S2: Histogram of flowering days in each treatment. Dashed lines represent the date of UAV measurements, Figure S3: The flow of plant area segmentation from each plot image. (a) Image of each plot, colored with a gradient according to the NDVI value; the plant area is within the green frame, the white mulching sheet area is within the blue frame, and the soil area is within the red frame. (b) Histogram of NDVI value within each color frame in (a); each color corresponds to the respective color of the frame. (c) Image masked with NDVI threshold of 0.15, Table S1: The description of accessions in each treatment. We used 198 accessions of soybean genetic resources, registered as the mini core collections in the National Institute of Agrobiological Sciences (NIAS) gene bank. Some accessions were removed due to the unavailability of phenotypic traits. NAs represent unavailable phenotypic data, while blanks represent data used in this study. Appendix S1: The details of kernels and single/multiple kernel models that were used in the analysis.

## Author Contributions

Y. Toda, Y. Ohmori, Y. Yamasaki, H. Takahashi, H. Takanashi, M. Tsuda, H. Tsujimoto, A. Kaga, M. Nakazono, T. Fujiwara, and H. Iwata acquired funding and conducted the field experiments. A. Kaga prepared the 198 accessions of soybean. H. Kajiya-Kanegae prepared the genome-wide marker data. K. Sakurai, Y. Toda, and H. Iwata designed the study and collected the multi-spectral image data. K. Sakurai analyzed the data and wrote a draft of the manuscript. H. Iwata supervised the study and edited the manuscript. All authors have reviewed and approve of the manuscript.

## Data Availability Statement

The datasets generated and analyzed in the present study are available from the “Sakuraikengo/TSMS_supple” repository in the GitHub, https://github.com/Sakuraikengo/TSMS_supple.

## Acknowledgments

We are grateful to the technical staff at the Arid Land Research Center, Tottori University, and Izumi Higashida. We would like to thank Kosuke Hamazaki for helping us with the ‘RAINBOWR’ package in R, Sawako Maruyama for making OMI of each band image, and Motoyuki Ishimori for fruitful discussions on how to use the MS image data.

