## Supplementary material for "Time-series Multi-spectral Imaging in Soybean for Improving Biomass and Genomic Prediction Accuracy": Figure S1, Figure S2, Figure S3, Table S1, Appendix S1: Supplementary.docx

**Table S1**	The description of accessions in each treatment. We used 198 accessions of soybean genetic resources, registered as the mini core collections in the National Institute of Agrobiological Sciences (NIAS) gene bank. Some accessions were removed due to the unavailability of phenotypic traits. NAs represent unavailable phenotypic data, while blanks represent data used in this study.

| Accession | Treatment C | Treatment W5 | Treatment W10 | Treatment D |
| --- | --- | --- | --- | --- |
| GmJMC002 |  |  |  | NA |
| GmJMC003 |  |  | NA |  |
| GmJMC004 |  |  |  |  |
| GmJMC005 |  |  |  |  |
| GmJMC007 |  |  |  |  |
| GmJMC008 |  |  |  |  |
| GmJMC009 |  |  |  |  |
| GmJMC013 |  |  |  |  |
| GmJMC016 |  |  |  |  |
| GmJMC017 |  |  |  |  |
| GmJMC021 |  |  |  |  |
| GmJMC023 |  |  |  |  |
| GmJMC025 |  |  |  |  |
| GmJMC026 |  |  |  |  |
| GmJMC028 |  |  |  |  |
| GmJMC030 |  |  |  |  |
| GmJMC031 |  |  |  |  |
| GmJMC032 |  |  |  |  |
| GmJMC033 |  |  |  |  |
| GmJMC034 | NA |  |  |  |
| GmJMC037 |  |  |  |  |
| GmJMC039 | NA |  |  |  |
| GmJMC040 |  |  |  |  |
| GmJMC041 |  |  |  |  |
| GmJMC043 |  |  |  |  |
| GmJMC044 |  |  |  |  |
| GmJMC047 |  |  |  |  |
| GmJMC049 | NA |  |  |  |
| GmJMC050 |  |  |  |  |
| GmJMC051 |  |  |  |  |
| GmJMC052 |  |  |  |  |
| GmJMC053 |  |  |  |  |
| GmJMC054 |  |  |  |  |
| GmJMC055 |  |  |  |  |
| GmJMC056 |  |  | NA |  |
| GmJMC057 |  |  |  |  |
| GmJMC058 |  |  |  |  |
| GmJMC059 |  |  |  |  |
| GmJMC060 |  |  |  |  |
| GmJMC061 |  |  |  |  |
| GmJMC062 |  |  |  |  |
| GmJMC063 |  |  |  |  |
| GmJMC064 |  |  |  |  |
| GmJMC065 |  |  |  |  |
| GmJMC067 |  |  |  |  |
| GmJMC068 |  |  |  |  |
| GmJMC069 |  |  |  |  |
| GmJMC076 |  |  |  |  |
| GmJMC077 |  |  |  |  |
| GmJMC078 |  |  |  |  |
| GmJMC079 |  |  |  |  |
| GmJMC080 |  |  |  |  |
| GmJMC081 |  |  | NA | NA |
| GmJMC082 |  |  |  |  |
| GmJMC085 |  |  |  |  |
| GmJMC088 |  | NA |  |  |
| GmJMC090 | NA | NA |  | NA |
| GmJMC091 |  |  |  |  |
| GmJMC092 |  |  |  |  |
| GmJMC093 |  |  |  |  |
| GmJMC095 |  |  |  |  |
| GmJMC096 |  |  |  |  |
| GmJMC097 |  |  | NA |  |
| GmJMC098 |  |  |  |  |
| GmJMC099 |  |  |  |  |
| GmJMC100 |  |  |  |  |
| GmJMC101 |  |  |  |  |
| GmJMC102 |  | NA |  |  |
| GmJMC104 |  |  | NA |  |
| GmJMC105 |  |  |  |  |
| GmJMC106 |  |  |  | NA |
| GmJMC110 |  |  |  |  |
| GmJMC111 |  |  |  |  |
| GmJMC112 |  |  | NA |  |
| GmJMC114 |  |  |  |  |
| GmJMC116 |  |  |  |  |
| GmJMC117 |  |  |  |  |
| GmJMC121 |  |  |  |  |
| GmJMC126 |  |  |  |  |
| GmJMC128 |  |  |  |  |
| GmJMC130 |  |  |  |  |
| GmJMC131 |  |  | NA |  |
| GmJMC133 |  |  |  |  |
| GmJMC137 |  |  |  |  |
| GmJMC139 |  |  |  |  |
| GmJMC145 |  |  |  |  |
| GmJMC149 |  |  |  |  |
| GmJMC158 |  |  |  | NA |
| GmJMC161 |  |  |  |  |
| GmJMC167 |  |  |  |  |
| GmJMC172 | NA |  |  |  |
| GmJMC177 |  |  |  |  |
| GmJMC179 |  |  |  |  |
| GmJMC180 |  |  |  |  |
| GmJMC184 |  |  |  |  |
| GmWMC001 |  |  |  |  |
| GmWMC006 |  |  |  |  |
| GmWMC010 |  |  |  |  |
| GmWMC011 |  |  |  |  |
| GmWMC012 |  |  |  |  |
| GmWMC014 |  |  |  | NA |
| GmWMC015 |  |  |  |  |
| GmWMC018 | NA |  |  |  |
| GmWMC019 |  |  |  |  |
| GmWMC020 | NA | NA | NA | NA |
| GmWMC022 |  |  |  |  |
| GmWMC024 |  |  |  |  |
| GmWMC027 |  |  |  |  |
| GmWMC029 |  |  |  |  |
| GmWMC035 |  |  |  |  |
| GmWMC036 |  |  |  |  |
| GmWMC038 | NA |  |  |  |
| GmWMC042 |  |  |  |  |
| GmWMC045 |  |  |  |  |
| GmWMC046 |  |  |  |  |
| GmWMC048 |  |  |  |  |
| GmWMC066 |  |  |  |  |
| GmWMC070 |  |  |  |  |
| GmWMC071 |  |  |  |  |
| GmWMC072 |  |  |  |  |
| GmWMC073 |  |  |  |  |
| GmWMC074 |  |  |  |  |
| GmWMC075 |  |  |  |  |
| GmWMC083 |  |  | NA |  |
| GmWMC084 | NA |  |  |  |
| GmWMC086 |  |  |  |  |
| GmWMC087 |  | NA |  |  |
| GmWMC089 |  |  |  |  |
| GmWMC094 |  |  |  | NA |
| GmWMC103 |  |  |  |  |
| GmWMC107 |  |  |  |  |
| GmWMC108 |  |  |  |  |
| GmWMC109 |  |  |  |  |
| GmWMC113 | NA | NA |  |  |
| GmWMC115 | NA |  |  |  |
| GmWMC118 |  |  |  |  |
| GmWMC119 |  |  |  |  |
| GmWMC120 |  |  |  |  |
| GmWMC122 |  |  | NA |  |
| GmWMC123 |  |  |  |  |
| GmWMC124 |  |  |  |  |
| GmWMC125 |  | NA |  |  |
| GmWMC127 |  |  |  |  |
| GmWMC129 | NA |  |  |  |
| GmWMC132 |  |  |  |  |
| GmWMC134 |  |  |  |  |
| GmWMC135 |  |  |  |  |
| GmWMC136 |  |  |  |  |
| GmWMC138 | NA |  |  | NA |
| GmWMC140 |  |  |  |  |
| GmWMC141 |  |  |  |  |
| GmWMC142 |  |  |  |  |
| GmWMC143 |  |  |  |  |
| GmWMC144 |  |  |  |  |
| GmWMC146 |  |  |  |  |
| GmWMC147 |  |  |  |  |
| GmWMC148 |  |  |  |  |
| GmWMC150 |  |  |  |  |
| GmWMC151 |  |  |  |  |
| GmWMC152 |  |  |  |  |
| GmWMC153 |  |  |  |  |
| GmWMC154 |  |  |  |  |
| GmWMC155 |  |  |  |  |
| GmWMC156 |  |  |  | NA |
| GmWMC157 |  |  |  |  |
| GmWMC159 | NA |  |  |  |
| GmWMC160 |  |  |  |  |
| GmWMC162 |  |  |  |  |
| GmWMC163 | NA |  |  |  |
| GmWMC164 |  |  |  |  |
| GmWMC165 | NA |  |  |  |
| GmWMC166 |  |  |  |  |
| GmWMC168 |  |  |  |  |
| GmWMC169 |  |  |  | NA |
| GmWMC170 |  |  |  |  |
| GmWMC171 |  |  |  | NA |
| GmWMC173 |  |  |  |  |
| GmWMC174 |  |  |  |  |
| GmWMC175 |  |  |  |  |
| GmWMC176 |  |  |  |  |
| GmWMC178 |  |  |  |  |
| GmWMC181 | NA |  |  |  |
| GmWMC182 |  |  |  |  |
| GmWMC183 |  |  | NA |  |
| GmWMC185 |  |  |  |  |
| GmWMC186 |  |  |  |  |
| GmWMC187 |  |  |  |  |
| GmWMC188 |  |  |  |  |
| GmWMC189 |  |  |  |  |
| GmWMC190 |  |  |  |  |
| GmWMC191 |  |  |  |  |
| GmWMC192 |  |  |  |  |
| Houjaku Kuwazu |  |  |  |  |
| Misuzudaizu | NA | NA | NA | NA |
| Norin2 | NA | NA | NA | NA |
| 5002T |  |  |  |  |
| C1329 |  |  |  |  |
| B01167 | NA | NA | NA | NA |

**Appendix S1**	The details of kernels and single/multiple kernel models that were used in the analysis.

**Kernels based on Relationship Matrices**

$\mathbf{V}_{each}$ was calculated for each combination of a growth stage (week) and a treatment. $\mathbf{V}_{each}$ was calculated as $\mathbf{S}_{each}\mathbf{S}_{each}^{T}$, where $\mathbf{S}_{each}$ is a $n\times t$ matrix of scaled phenotypic values of VIs (t = 5) for each genotype. $\mathbf{V}_{all}$ was calculated for each growth stage over all treatments. $\mathbf{V}_{all}$ was calculated as $\mathbf{S}_{all}\mathbf{S}_{all}^{T}$, where $\mathbf{S}_{all}$ is a $nm\times t$ matrix of scaled phenotypic values of VIs for each plot. $nm$ indicates the number of plot over all treatments. A kernel based on the field heterogeneity relationship matrix ($\mathbf{FH}$) was defined based on the Euclidean distances between plots based on the positions in the field. Plots belonging to different rows were assumed to be independent; thus, the Euclidean distance was set to infinity in this case. The Euclidean distances were converted to a Gaussian kernel, where the ($i$, $j$)-th element of said kernel is defined as:

$$f_{ij}=\text{exp}\left( -hd_{ij}^{2} \right)$$

where $d_{ij}^{2}$ is the squared distance between plots and $h$ is the reciprocal of the median of squared distances between all combinations of all plots.

**Single/Multiple Kernel Models within Each Treatment**

**The Additive genomic relationship kernel model (*G* model)**

This model predicts the phenotypic values of AGB using only genome-wide marker data. As a basis for comparison to assess the advantage of VIs, the following single-kernel regression model was built using the additive genomic relationship kernel ($\mathbf{G}$):

$$\begin{aligned} y_{il}=b_{l}+u_{i}+\varepsilon_{i}\#\left( S1 \right) \end{aligned}$$

where $y_{il}$ is the phenotypic value of AGB for genotype $i$ $\left( i=1,...,n \right)$. $b_{l}$ the fixed effect of flowering $\left( l=1,2 \right)$, $u_{i}$ the genetic random effect, and $\varepsilon_{i}$ the residual random effect. The vector $\mathbf{u}=\left( u_{1},...,u_{n} \right)^{T}$ follows the multi-variate normal distribution $MVN\left( \mathbf{0},\mathbf{G}\sigma_{g}^{2} \right)$, where $\mathbf{G}$ is the additive numerator relationship kernel and $\sigma_{g}^{2}$ is the additive genetic variance. The vector $\boldsymbol{\varepsilon}=\left( \varepsilon_{1},...,\varepsilon_{n} \right)^{T}$ follows the multi-variate normal distribution $MVN\left( \mathbf{0},\mathbf{I}\sigma_{e}^{2} \right)$, where $\mathbf{I}$ is the identical matrix and $\sigma_{e}^{2}$ is the residual variance.

**The VI relationship kernel model (**$\boldsymbol{V}_{\boldsymbol{each}}$ **model)**

To evaluate the usefulness of VIs in predicting the phenotypic values of AGB at each irrigation level, the following single-kernel model was built based on $\mathbf{V}_{each}$:

$$\begin{aligned} y_{il}=b_{l}+v_{i}+\varepsilon_{i}\#\left( S2 \right) \end{aligned}$$

where $y_{i}$, $b_{l}$, and $\varepsilon_{i}$ are defined as above in the model of Equation S1, and $v_{i}$ is the random effect of the VIs. The vector $\mathbf{v}=\left( v_{1},...,v_{n} \right)^{T}$ follows the multi-variate normal distribution $MVN\left( \mathbf{0},\mathbf{V}_{each}\sigma_{v}^{2} \right)$, where $\mathbf{V}_{each}$ is the VIs relationship kernel for each treatment and $\sigma_{v}^{2}$ is the VIs variance.

**Model with** $\mathbf{G}$ **and** $\mathbf{V}_{\boldsymbol{each}}$ **kernels (**$\boldsymbol{G}\mathbf{+}\boldsymbol{V}_{\boldsymbol{each}}$ **model)**

By using two types of kernels, the additive genomic relationship kernel ($\mathbf{G}$) and VIs relationship kernel ($\mathbf{V}_{each}$), we can make predictions using both genome and VIs data. This multi-kernel model was built using ($\mathbf{G}$) and ($\mathbf{V}_{each}$):

$$\begin{aligned} y_{il}=b_{l}+u_{i}+v_{i}+\varepsilon_{i}\#\left( S3 \right) \end{aligned}$$

Here, $y_{il}$, $b_{l}$, $u_{i}$, and $\varepsilon_{i}$ are defined as in the model of Equation S1, while $v_{i}$ is defined as in the model of Equation S2.

**Model with** $\mathbf{G}$ **kernel and field heterogeneity relationship kernel (**$\boldsymbol{G}\mathbf{+}\boldsymbol{Field}$ **model)**

To confirm that there was a field heterogeneity resulted in the distances among plots or not in the experimental field. This multi-kernel model was built using additive genomic relationship kernel ($\mathbf{G}$) and the field heterogeneity relationship kernel ($\mathbf{FH}$):

$$\begin{aligned} y_{il}=b_{l}+u_{i}+f_{i}+\varepsilon_{i}\#\left( S4 \right) \end{aligned}$$

Here, $y_{il}$, $b_{l}$, $u_{i}$, and $\varepsilon_{i}$ are defined as in the model of Equation S1, and $f_{i}$ is the random effect reflecting the among-plot distances. The vector $\mathbf{f}=\left( f_{1},...,f_{n} \right)^{T}$ follows the multi-variate normal distribution $MVN\left( \mathbf{0},\mathbf{FH}\sigma_{f}^{2} \right)$, where $\mathbf{FH}$ is the field heterogeneity relationship matrix and $\sigma_{f}^{2}$ is the variance of field heterogeneity based on the among-plot distances.

**Single/Multiple Kernel Models over All Treatments**

**Model with** $\mathbf{G}$ **kernel and treatment and genotype-by-treatment interaction effects (**$\boldsymbol{G}\mathbf{+}\boldsymbol{E}\mathbf{+}\boldsymbol{G}\boldsymbol{\times}\boldsymbol{E}$ **model)**

This model assumes that genome-wide marker data and treatment information are available. The effects of treatment and genotype-by-environment interaction were added to the $G$ model :

$$\begin{aligned} y_{ijl}=E_{j}+b_{l}+u_{i}+ge_{ij}+\varepsilon_{ij}\#\left( S5 \right) \end{aligned}$$

Here, $u_{i}$ $\left( i=1,...,n \right)$is defined as above in the model of Equation S1, $y_{ij}$ is the phenotypic trait value of AGB for genotype $i$ in treatment $j$, $E_{j}$ is the fixed effect for treatment $j$ $\left( j=1,...,m \right)$, $b_{l}$ is the fixed effect of flowering or not $\left( l=1,2 \right)$, $ge_{ij}$ is the genotype-by-environment interaction random effect, and $\varepsilon_{ij}$ is the residual random effect. $\mathbf{ge}=\left( ge_{11},...,ge_{nm} \right)^{T}$ follows multi-normal distribution $MVN\left( \mathbf{0},\left( {\mathbf{Z}_{\mathbf{g}}\mathbf{G}\mathbf{Z}_{\mathbf{g}}}^{T} \right)\circ\left( {\mathbf{Z}_{\mathbf{E}}\mathbf{Z}_{\mathbf{E}}}^{T} \right)\sigma_{ge}^{2} \right)$, where $\mathbf{Z}_{\mathbf{g}}$ and $\mathbf{Z}_{\mathbf{E}}$ are incidence matrices for genotypes and treatments, and $\sigma_{ge}^{2}$ is the variance component for the effect of genotype-by-environment interaction [1][2][3]. $\boldsymbol{\varepsilon}=\left( \varepsilon_{11},...,\varepsilon_{nm} \right)^{T}$ follows the multi-normal distribution $MVN\left( \mathbf{0},\mathbf{I}\sigma_{e}^{2} \right)$, where $\mathbf{I}$ is the identical matrix and, $\sigma_{e}^{2}$ is the residual variance.

**Model with** $\mathbf{G}$ **and** $\mathbf{V}_{\boldsymbol{all}}$ **kernels and the fixed effect of treatment (**$\boldsymbol{G}\mathbf{+}\boldsymbol{E}\mathbf{+}\boldsymbol{V}_{\boldsymbol{all}}$ **model)**

To model the $G\times E$ effect using the VIs relationship kernel ($\mathbf{V}_{all}$), this multi-kernel model includes genome-wide marker data, treatment information, and phenotypic traits of VIs:

$$\begin{aligned} y_{ijl}=E_{j}+b_{l}+u_{i}+v_{ij}+\varepsilon_{ij}\#\left( S6 \right) \end{aligned}$$

Here, $y_{ijl}$, $E_{j}$, $b_{l}$, $u_{i}$, and $\varepsilon_{ij}$ are defined as above in model Equation S5. $v_{ij}$ is the random effect of the VIs. The vector $\mathbf{v}=\left( v_{11},...,v_{nm} \right)^{T}$ follows the multi-variate normal distribution $\mathbf{v}\sim MVN\left( \mathbf{0},\mathbf{V}_{all}\sigma_{v}^{2} \right)$, where $\mathbf{V}_{all}$ is the VIs relationship kernel over all treatment and $\sigma_{v}^{2}$ is the VIs variance.

**Model with** $\mathbf{G}$ **and** $\mathbf{V}_{\boldsymbol{all}}$ **kernels (**$\boldsymbol{G}\mathbf{+}\boldsymbol{V}_{\boldsymbol{all}}$ **model)**

This model evaluates whether the VIs can capture the macro-environmental variability, i.e., the differences in irrigation levels. This model predicts the phenotypic values of AGB from marker information and the phenotypic data of VIs without treatment information. This model was nearly the same for Equation S3 :

$$\begin{aligned} y_{ijl}=b_{l}+u_{i}+v_{ij}+\varepsilon_{ij}\#\left( S7 \right) \end{aligned}$$

Here, $y_{ij}$, $b_{l}$, $u_{i}$, and $\varepsilon_{ij}$ are defined as above in Equation S5. $v_{ij}$ is defined as above in Equation S6.

**Model with** $\mathbf{V}_{\boldsymbol{all}}$ **kernels (**$\boldsymbol{V}_{\boldsymbol{all}}$ **model)**

This model was built using only VIs phenotypic data, considering the that neither genome-wide marker data nor treatment information were available. This model was almost the same for Equation S2 :

$$\begin{aligned} y_{ijl}=b_{l}+v_{ij}+\varepsilon_{ij}\#\left( S8 \right) \end{aligned}$$

Here, $y_{ijl}$, $b_{l}$, and $\varepsilon_{ij}$ are defined as above in model Equation S5. $v_{ij}$ is defined as above in model Equation S6.
