## Supplementary figures and images for "Time-series Multi-spectral Imaging in Soybean for Improving Biomass and Genomic Prediction Accuracy"

### FigureS1.jpg

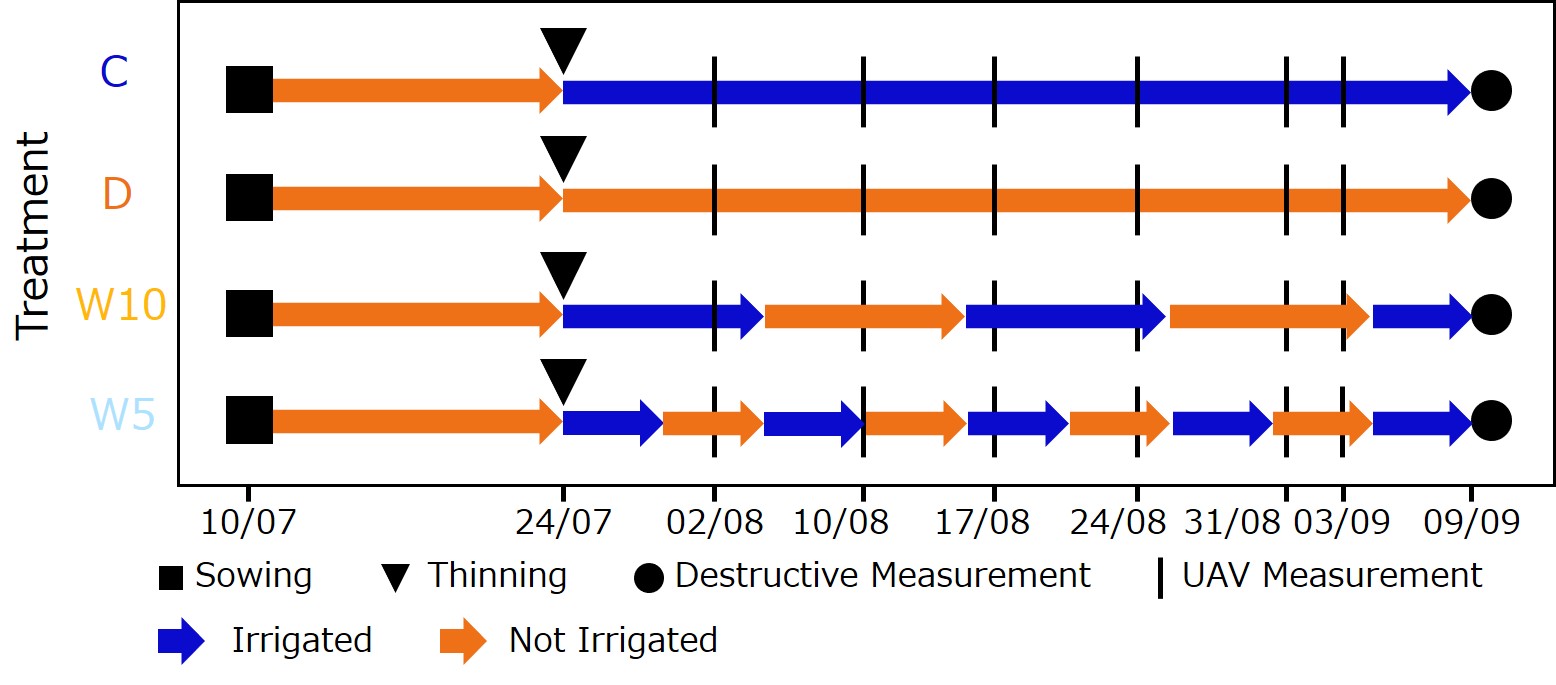

### FigureS2.jpg

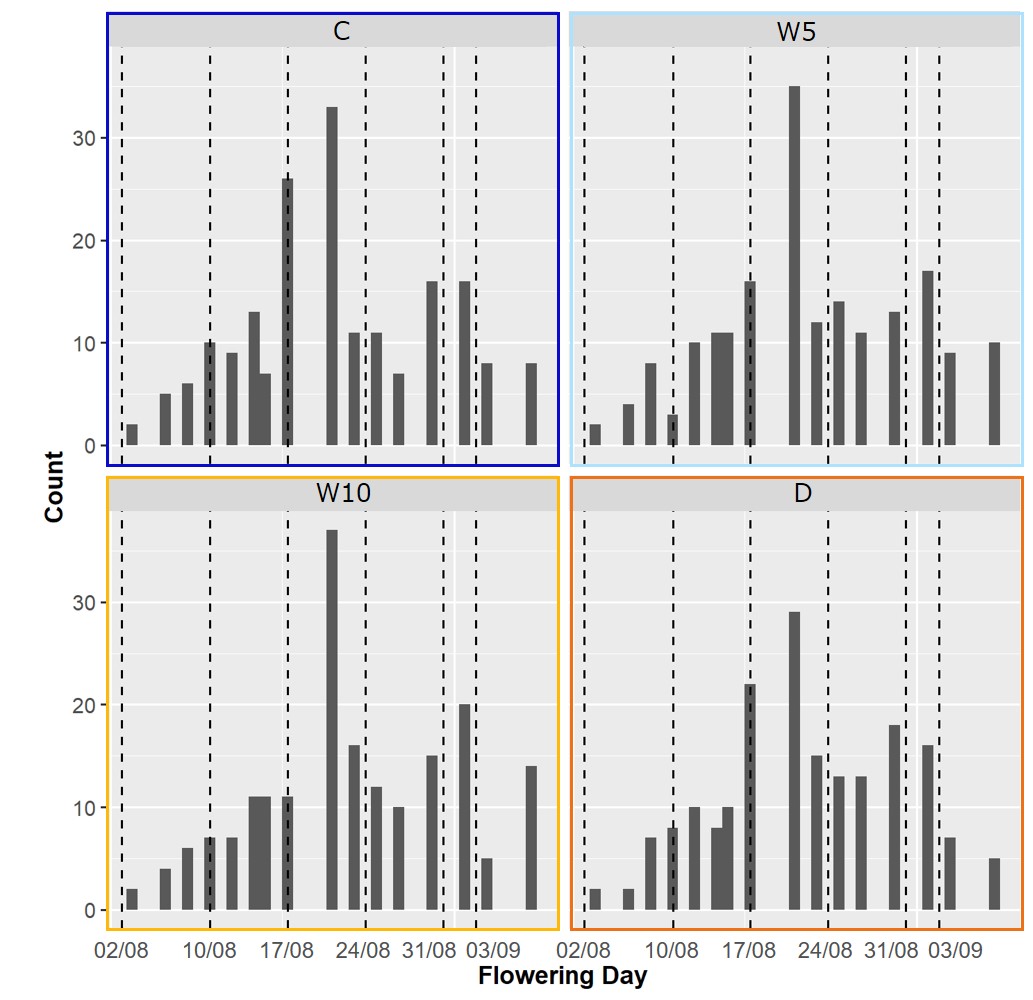

### FigureS3.jpg

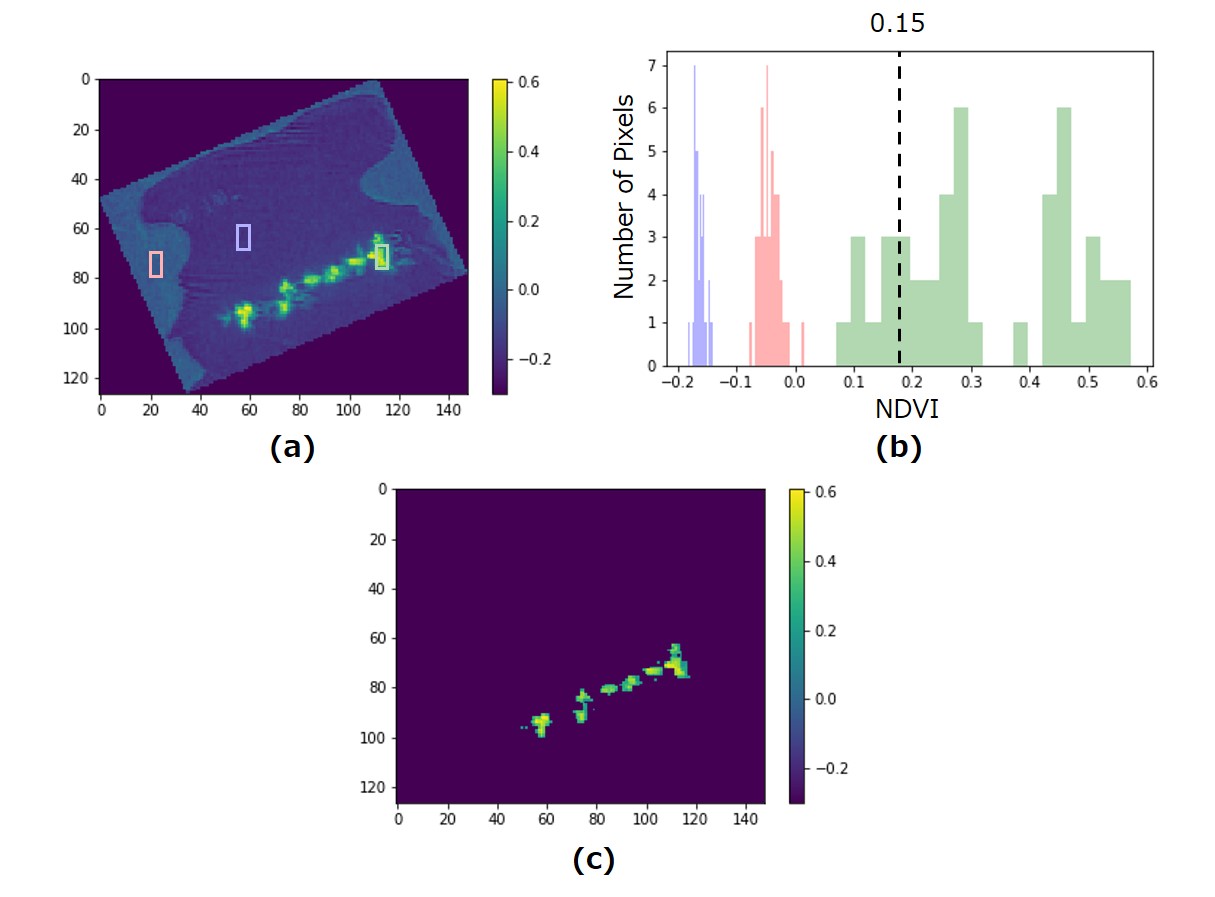
